# Respiratory syncytial virus nonstructural protein 1 inhibits production of cytokines and chemokines by differentiated primary nasal epithelial cells cultured at air-liquid interface

**DOI:** 10.1101/2025.10.20.683500

**Authors:** Rosanne W Koutstaal, Amadeo Munoz Garcia, Nadine Ebert, Manon F Licheri, Anke J Lakerveld, Hendrik-Jan Hamstra, Anne T Gelderloos, Jørgen de Jonge, Cécile ACM van Els, Volker Thiel, Ronald Dijkman, Puck B van Kasteren

## Abstract

Respiratory syncytial virus (RSV) is a major cause of severe respiratory infections in children and older adults. Several vaccines have recently been approved, which are all based on the RSV fusion (F) protein. Among the strategies to induce a broader immune response is the development of live-attenuated RSV vaccines, for example by disabling nonstructural protein 1 (NS1). RSV NS1 has been found to modulate the host immune response by repression of interferon (IFN) production. However, functional characterization of this protein was mainly performed in immortalized cell lines, which do not necessarily represent the *in vivo* situation.

Here, we assessed the effect of NS1 mutations on replication and host responses in differentiated primary human nasal epithelial cells (HNEC) cultured at air-liquid interface (ALI), which mimic several physiological properties of the human airways. Using a newly developed yeast-based RSV reverse genetics system, inactivating mutations in the NS1 gene were introduced. These mutations did not affect viral replication in Vero and A549 cells. However, in HNEC we observed a delayed replication of the NS1 mutants compared to the WT virus. Bulk RNA sequencing data from early post infection revealed a stronger antiviral signature in HNEC that were infected with NS1 mutants compared to WT, characterized by upregulation of IFN-β and IFN-λ_1/2/3_ and chemokines CXCL11 and CCL5. This signature was confirmed by analysis of the cytokines that were released into the HNEC basal medium. Finally, the basal medium containing these cytokines was used for an indirect immune cell migration assay. In this set-up, both WT and NS1 mutant viruses induce migration of mainly neutrophils.

In conclusion, this study shows that RSV NS1 supports viral replication not only via inhibition of the production of IFNs, but also by reducing early chemokine production and secretion by epithelial cells. Furthermore, our data highlight the suitability of the ALI transwell model for preclinical assessment of live-attenuated vaccine candidates.

## Introduction

Human respiratory syncytial virus (RSV) infection is one of the leading global causes of severe respiratory disease in children and older adults. Among the available prevention strategies are prophylactic monoclonal antibodies, two subunit vaccines and an mRNA vaccine [1-5]. A limitation of the currently approved vaccines is that all of them are focused on RSV F-mediated immunity. The use of a single antigen increases the risk for the emergence of escape mutants. In addition, F-based vaccines have been associated with vaccine-enhanced disease in naïve children [6-9]. To broaden the vaccine-induced immune response, it would be of value to include additional antigens in RSV vaccines. One of the ways to achieve such a broad response is the use of live-attenuated vaccines, similar to successful vaccines against for example influenza, measles, mumps, polio, and rotavirus [10, 11]. For attenuation, the viral genome is adapted in such a way that the virus loses its virulence, while retaining its replicative capacity and immunogenicity. To this end, viruses can be rationally edited using recombinant techniques or passively altered via serial passaging [12]. Live-attenuated RSV is expected to induce both innate immunity and durable local and systemic immunity [13-15]. Moreover, live-attenuated vaccines have not been associated with vaccine-enhanced RSV disease [16].

Research into live-attenuated RSV vaccine candidates has been ongoing for decades [12]. However, most of these fail to pass the phase I/II stage of clinical development, highlighting the need for improved preclinical models for the evaluation of promising candidates. Preclinical development currently relies mainly on (small) animal models, which often do not faithfully recapitulate the pathogenesis observed for RSV in humans [17-19], and *in vitro* culture models of immortalized cell lines. Firstly, these cells often contain genetic abnormalities – which further accumulate during passaging – of which the impact on (antiviral) host responses is unclear. Secondly, these cell lines often lack the cellular receptors that are relevant for RSV binding and entry, like the fractalkine receptor CX3CR1 [20, 21]. In the upper respiratory tract, where the primary infection takes place, multiple cell types are present that each have their own function during infection. For example, goblet cells excrete mucus, which creates a protective barrier against pathogens and contains numerous types of antimicrobial peptides [22]. This mucus is transported over the surface of the epithelium by ciliated cells – which are the main target cells for RSV. Differentiated primary airway epithelial cells (AEC) grown at air-liquid interface (ALI) provide a suitable infection model for RSV that contains both of these cell types [23-25], forming a bridge between non-human animal models, immortalized cell lines and humans.

Several candidate live-attenuated vaccine strains contain alterations in the genes that encode for RSV nonstructural protein (NS) 1 and 2 [26, 27]. These proteins were found to be involved in immune evasion, which makes them interesting targets for attenuation. RSV mutant strains without functional NS1 and/or NS2 showed reduced replication both *in vitro* and *in vivo*, while retaining their immunogenic capacities [28, 17, 18, 29]. Schlender *et al*. demonstrated that bovine RSV NS1 and NS2 cooperatively inhibit the type I IFN response of the infected host cell [30]. Of these two proteins, NS1 seems to be the strongest inhibitor, while the presence of NS2 enhances this effect. Human RSV NS1 was found to interfere with both type I and type III interferons in (immortalized) epithelial cells as well as in primary macrophages [31]. The (molecular) mechanisms behind NS1 and NS2-mediated interference with IFN have been described in a review by Van Royen *et al*. [32]. In addition to the inhibition of IFN responses in airway epithelial cells, RSV NS1 has been found to interfere with the activation of adaptive immune responses, like dendritic cell (DC) maturation and T cell homing to the airway mucosa [33-35]. These findings could – at least in part – explain the lack of a durable immune response to RSV. However, the precise role of NS1 in the interaction between the airway epithelium and immune cells in the context of an RSV infection remains to be elucidated. Importantly, most of the abovementioned studies into the function of the RSV nonstructural proteins were performed in immortalized cell lines and based on either ectopic expression or complete NS1 deletion mutants.

In the current study, the ALI transwell model is used to study the effect of mutations in the NS1 α3 helix to obtain insights into its role in immune evasion in a more physiologically relevant model. In ectopic expression experiments, these rationally designed mutations were previously demonstrated by Chatterjee *et al*. to result in functional impairment of the NS1 protein while not disrupting its structural integrity, showing the importance of the α3 helix for immune evasion [36]. Since primary AEC cultured at ALI have a functional IFN system, we hypothesized that the NS1 mutants would show reduced replication in this model, accompanied by an increased production of type I and III IFNs. In order to investigate this, the growth kinetics of wild-type (WT) and NS1 mutant RSV were first assessed on immortalized cell lines (Vero/A549) and primary human nasal epithelial cells (HNEC) at ALI. In addition, host responses to these viruses were assessed by bulk RNA sequencing and determination of cytokine/chemokine levels in the basal medium of WT or mutant RSV-infected ALI cultures. Finally, the effect of secreted mediators on immune cell migration was investigated. Taken together, our data provide insights into the effect of NS1 on the innate immune response in a physiologically relevant human respiratory epithelial cell culture model, showing the potential value of this model for preclinical assessment of live-attenuated RSV vaccine candidates.

## Materials & Methods

### Ethics

This study was conducted according to the principles described in the Declaration of Helsinki, and for the collection of blood samples and subsequent analyses, all donors provided written informed consent. Blood samples were processed anonymously and the research goal, primary cell isolation, required no review by an accredited Medical Research Ethics Committee (MREC), as determined by the Dutch Central Committee on Research involving human subjects (CCMO). Primary epithelial cells were procured from Epithelix. Epithelial cells were processed anonymously by Epithelix partner collect centers, and written informed consent was obtained from all donors or relatives before collection.

### Cell lines

Vero cells (RRID: CVCL_0059, ATCC CCL-81) were cultured in Dulbecco’s modified Eagle’s medium (DMEM; Gibco, 41965-062) supplemented with 5% heat-inactivated (hi) fetal bovine serum (FBS) (HyClone, SH30071.03) and 1% penicillin/streptomycin/glutamine (P/S/G) (cat#10378-016, Gibco). BSR-T7/5 cells (RRID: CVCL_RW96) were cultured in DMEM + 5% FBS + 1% P/S/G and 1mg/mL G418 sulphate. A549 cells (RRID:CVCL_LI35, ATCC CCL-185) were cultured in RPMI medium 1640 (Gibco, 52400-025) + 5% FBS + 1% P/S/G. HEp-2/HeLa cells (RRID:CVCL_1906, ATCC CCL-23) were cultured in Minimal essential medium (MEM) (Gibco; 31095-029) supplemented with 10% FBS and 1% P/S/G.

### Reverse genetics

Full-length clones of wild-type and mutant HRSV/A/NLD/25147X/1998 (GenBank: FJ948820) were constructed by transformation-associated recombination (TAR) cloning in yeast, as described previously [37]. Briefly, 7-8 overlapping PCR products encompassing the viral genome, accessory sequences, and vector pCC1BAC-His3 [38] were transformed into yeast strain VL6-48N [39] using the lithium acetate/SS carrier DNA/PEG method [40]. The viral genome is flanked by an upstream T7 promotor and hammerhead ribozyme sequence and a downstream hepatitis delta virus ribozyme, and mutations (Y125A or L132A/L133A) were introduced via mismatched oligonucleotides. Transformed yeast was grown on SD-His agar plates until colonies appeared (2-4 days) and individual yeast colonies were subsequently expanded by growing in SD-His broth for 18 h at 30°C at 220 rpm. Assembled full-length clones were isolated from yeast using the Plasmid DNA purification kit (Machery-Nagel, Germany) and Zymolyase (Sigma-Aldrich, USA) and amplified in Transformax EPI300 *E. coli* (C300C105, LGC Biosearch Technologies, Hoddesdon, UK) using CopyControl induction solution (CCIS125, LGC Biosearch Technologies). Genome sequences were verified by whole plasmid nanopore sequencing (Plasmidsaurus, USA), where remaining uncertainties were assessed by Sanger sequencing.

### Virus rescue

Recombinant viruses were rescued by transfecting the full-length clones (4.8 μg) together with helper plasmids pcDNA3.1-L (0.4 μg), pcDNA3.1-N (1.6 μg), pcDNA3.1-P (1.2 μg), pcDNA3.1-M2-1 (1 µg) and pCAGGS-T7 (1 μg) into BSR-T7/5 cells using Lipofectamine LTX with Plus reagent (Thermo Fisher Scientific, USA) according to the manufacturer’s instruction in a 6-well plate format. At 3 days after transfection, virus-containing supernatants were collected and passaged 3-4 times at MOI ≤0.02 for 3-7 days on Vero cells in medium containing 2% FCS. Virus-containing supernatants were finally harvested, cleared by centrifugation, supplemented with sucrose at a final concentration of 10%, aliquoted, snap-frozen and stored at -80°C as master stocks.

### Virus stocks and quantification

Sucrose-purified virus stocks were produced by infecting HEp-2/HeLa cells with an MOI of 0.02 for 3 days using the passage 4 master stocks. Virus stocks were purified between layers of 10 and 50% sucrose by ultracentrifugation, washed with plain RPMI and concentrated on 100 kDa filters (Millipore cat. no. UFC910024) by centrifugation. The virus was washed from the filters in phenol red-free RPMI (Gibco cat. no. 11835-030) supplemented with 10% sucrose and subsequently snap-frozen and stored at -80°C. NS1 and NS2 genes were Sanger sequenced to confirm presence of the mutations.

Virus quantification was performed by 50% tissue culture infectious dose assay on Vero cells (TCID50_Vero_). In brief, samples were serially diluted 1:8 in DMEM (Gibco, 41965-062), supplemented with 100 units/mL Pen and 0.1 mg/mL Strep (Sigma, #P4333) and 2% FCS (HyClone, SH30071.03). Virus dilutions were added in quadruplicate to a confluent monolayer of Vero CCL-81 cells, and incubated at 37°C for 6 days. Subsequently, each well was scored either negative or positive for cytopathic effects, and viral load was determined using the Spearman & Kärber method [41].

### ALI cultures

Primary human nasal epithelial cells (HNEC) from 3 healthy donors (M/35y, F,/41y, M/50y; EP51AB, Epithelix, Switzerland) were expanded in complete PneumaCult Ex Plus medium (Stemcell Technologies, #05040, Canada) until the cells reached log phase. Cells were then harvested using trypsin (Lonza, CC-5012), which was inactivated using trypsin neutralizing solution (Lonza, CC-5002). Passage 1 cells were frozen in 78% Ham’s nutrient mix F12 (Merck/Sigma, 51651C) with 10% FCS, 2% 1.5M Hepes [pH 7.2], and 10% DMSO, and stored at -135°C until further use.

Stored cells were thawed in Complete PneumaCult Ex Plus medium in a T75 culture flask and expanded until log phase was reached, replacing the medium every 1-2 days. The cells were harvested using trypsin as described above and seeded at a density of 8×10^4^ cells/well on the apical side of transwell inserts with a PET membrane with pore size 0.4 μm and diameter 6.5mm (Corning, #CLS3470, New York, USA) in 100 μL PneumaCult Ex Plus medium. In addition, 650 μL culture medium was added to the basolateral compartment. After 48-72 hours, confluent monolayers were airlifted by aspirating the media from both compartments and adding 500 μL complete PneumaCult ALI medium (Stemcell Technologies, #05001) supplemented with 100 units/mL Penicillin and 0.1 mg/mL Streptomycin (Pen/Strep, Sigma, P4333-100mL), 0.48 μg/mL Hydrocortisone (Stemcell Technologies, #07926) and 4 μg/mL Heparin solution (Stemcell Technologies, #07980), to the basolateral compartment only. Medium was replaced every 2-3 days (500 μL/well for 2 days, 650 μL/well for 3 days of culture). From week 2 post airlift, the apical side of the cells was washed 1-2 times per week using 200 μL PBS (Gibco, #20012-068) to remove excess mucus. Cells were kept in a humidified incubator at 37°C and 5% CO2.

### Viral infection and sampling procedures

#### Infection of primary HNEC

Differentiated HNEC were used for infection between 5-7 weeks post airlifting. Three days before infection, the basal cell culture medium was replaced with complete PneumaCult ALI medium without heparin and hydrocortisone (ALI infection medium). Directly before infection, cells were washed apically using 200 μL/well HBSS (Gibco, #14025092) for 10 min at 37°C, and the basal medium was replaced with 500 µL infection medium/well. Sucrose-purified virus was diluted to 5×10^5^ TCID50/mL in HBSS. Cells were infected apically in triplicate using 200 μL inoculum, corresponding to 1×10^5^ TCID50 per well, and incubated at 37°C and 5% CO2 for 1 h, after which the inoculum was aspirated. Cells were immediately washed thrice with 200 μL HBSS.

For virus quantification, apical washes were performed at 2, 24, 48, 72 and 96 hours post infection (hpi) by adding 200μL HBSS and incubating 15 min at 37°C. 140 μL of the samples was snap-frozen using 100% EtOH in dry ice and stored at -80°C until further analysis. The other part of the samples was stored in MagNA pure External Lysis Buffer (Roche, 06374913001) for subsequent RNA isolation. In parallel with the apical washes, 50 μL basolateral medium was obtained from each well for cytokine analysis. These samples were also snap-frozen and stored at -80°C. For the bulk RNA sequencing experiment, the same infection procedure was used. At 20 hpi, whole cell lysates were obtained by adding 90 μL MagNA pure External Lysis Buffer to each transwell, which was incubated at RT for 10 min. Samples were resuspended in 60 uL HBSS and stored at -80°C until further analysis.

#### Infection of Vero/A549

Confluent monolayers of Vero and A549 cells were infected in triplicate in a 24-well plate at an MOI of 0.02. Cell culture supernatant samples were taken at 2, 24, 48, 72 and 96 hpi. Of these samples, one part was stored in MagNA pure External Lysis Buffer, and the other part was snap-frozen and stored at -80°C.

### RNA isolation

120 μL sample in lysis buffer was added to 50 μL AMPure XP Reagent (Beckman Coulter, A63881) in a 96-well PCR plate and incubated for 20 minutes to allow RNA binding to the beads. The beads were pulled down using a DynaMag™-96 Side Skirted Magnet (Invitrogen, 12027) and washed 3x using 70% EtOH (100% EtOH (VWR, 20.816) mixed with ultrapure water). RNA was eluted by resuspending the beads in nuclease free water (Ambion, AM9937) and samples were stored at -80°C until further use.

### RT-qPCR

The following primers were used to quantify RSV RNA in apical wash samples: RSV-A fw: TGAACAACCCAAAAGCATCA, RSV-A rev: CCTAGGCCAGCAGCATTG and probe: AATTTCCTCACTTCTCTAGTGTAGTATTGGG with a FAM reporter and BHQ1 quencher. Viral RNA was reverse-transcribed and amplified in one reaction using the TaqMan™ Fast Virus 1-Step Master Mix for qPCR (Applied Biosystems, 4444432), using 5 μL RNA per reaction in a total reaction volume of 20 μL. RT-qPCR was performed on the StepOnePlus System (Applied Biosystems). Serially diluted virus stocks with a known concentration of viral particles were used as a reference standard.

### Bulk RNA sequencing library preparation

The NEBNext Ultra II Directional RNA Library Prep Kit for Illumina was used to process the sample(s). The sample preparation was performed according to the protocol “NEBNext Ultra II Directional RNA Library Prep Kit for Illumina” (NEB #E7760S/L). The sequencing was performed using an Illumina NovaSeq 6000 sequencer.

### RNA bulk sequence data preprocessing

Adapter sequences (TruSeq adapters) and low-quality bases were trimmed using fastp v.0.23.2 [42]. Reads of which ≥40% of their bases had an average phred score below Q15 or had more than 5 unknown bases, were excluded. Reads shorter than 15 pb were excluded (both pairs in paired-end reads). Filtered reads were mapped to the human GRCh38.p13 reference genome using STAR2 v2.7.10 [43]. Count table was generated by HTSeq v2.0.2 [44]. A median of 61 M paired reads were filtered of which a median of 59 M were mapped.

### Downstream analysis of RNA bulk sequencing

For each mutation, we performed a differential expression gene (DEG) analysis using the R package DESeq2 [45]. We performed the standard DEG analysis that is wrapped in the single function DESeq(), therefore a Wald test was used. For each mutation, a comparison to the wild-type was performed. In the design formula, we accounted for paired-samples controlling for donor-level confounders. We pre-filtered the genes that at least have 10 counts in at least 3 samples as specified in the DESeq2 standards for RNA-bulk sequencing analysis. Gene set enrichment analysis (GSEA) was performed using the ranked gene lists of the significantly altered genes (adjusted p value < 0.05) of each condition. Genes were ranked based on logFC. The GSEA was performed using the clusterProfiler R package and the gene ontology (GO) database with the biological process annotation [46-48].

### Multiplex cytokine analysis

Cytokine concentrations were determined using LEGENDplex™ custom human panel 1805 (Biolegend) according to the manufacturer’s instructions, except for the use of a reduced volume per reaction. This panel contained the following cytokines: IFN-λ1, IFN-λ2/3, IL-8, IFN-α2, CXCL10, CCL2, CCL5, CXCL11, CXCL9, IL-6, IL-18 and IFN-β. The data was analysed using the Qognit software provided by Biolegend. All concentrations below the detection limit were set to ([detection limit]/2). To calculate fold changes in cytokine levels compared to WT, all individual values were divided by the mean of the WT condition of the respective timepoint.

### Immune cell migration assay

Whole blood from healthy donors was collected in heparin tubes. Blood was diluted 1:1 in PBS. A double density gradient was prepared by layering 10 mL Ficoll Paque Plus (Cytiva, 17-1440-02) onto 12 mL Histopaque-1119 (Sigma, 11191) in a 50 mL tube. On top, 28 mL diluted blood was added. The gradients were centrifuged 20 min at 396 xg, acc. 5, dec. 1 at RT. After centrifugation, the plasma was discarded and the PBMC and granulocyte fractions were collected separately. Cells were washed 2x in RPMI + 0,1% human serum albumin (HSA; Albuman 200 g/L, Sanquin, NL) and 1% P/S/G. On the granulocytes, a hyperosmotic shock was performed to lyse the erythrocytes, by adding 9 mL ultrapure water for 30 seconds, after which 1 mL 10x PBS was added to stop the reaction.

Both PBMCs and granulocytes were counted and diluted to 6,25×10^6^ cells/mL in RPMI + 0,1% HSA + P/S/G. PBMCs and granulocytes were mixed in a 1:1 ratio (approaching the *in vivo* proportions [49]), after which 80 μL (5×10^5^ cells/well) was added to the apical compartment of a 96-well transwell plate with a 3.0 μm pore-size membrane (Corning, 3385). To the basal compartment of the 96-well plate, basolateral samples from RSV-infected HNEC cultures were added at a 1:10 dilution in ALI infection medium, using 100 μL per well. The basal samples contained pooled medium from 3 different HNEC donors (from a single infection experiment), that had been infected with RSV for 24 or 48 h. The 96-wells plate containing the immune cells and basal medium samples was incubated for 1 h at 37°C to allow cells to migrate from the apical to the basal side.

After incubation, the cells from both the apical and the basal compartment were harvested. Both compartments were incubated with StemPro Accutase (Gibco, A1110501) for 15 min to ensure all cells had detached from the plate. The cells from the basal compartment were stained with CD45 BUV395 (1:100; BD, 563792) to label all migrated cells, after which the cells from the apical and basal compartments were combined. Subsequently, all samples were stained with the following panel: CD4 PerCP Cy5.5 (1:50; Biolegend, 399530), CD66b BB515 (1:100; BD, 564679), CD3 AF700 (1:40; BD, 659119), Live-dead eFluor780 (1:1000; Invitrogen, 65-0865-14), CD16 BV510 (1:50; Biolegend, 302048), CD56 BV711 (1:20; Biolegend, 318336), CD8 BV786 (1:100; BD, 563823), CD14 PE (1:100; Biolegend, 301806), CD19 PE-CF594 (1:50; Biolegend, 302252), CD11c PE-Cy7 (1:160; Biolegend, 301608), CD11b BUV737 (1:100; BD, 612800). After staining, the cells were fixed using 4% formaldehyde (VWR, #9713.5000) and measured using a BD FACS Fortessa. Data analysis was performed using FlowJo V10_8_1. The gating strategy can be found in **Supplementary Figure 1a-b**.

## Results

### RSV NS1 mutations are stable during in vitro infection of epithelial cells

Using a novel yeast-based reverse genetics system for RSV - building on previous work by Thi Nhu Thao *et al*. - we inserted two sets of mutations in a 1998 RSV/A strain [37]. Amino acid substitutions Y125A and L132/133A were achieved by mutating 3 or 6 nucleotides, respectively, as depicted in **Figure 1a**. These mutations are not naturally occurring, but have been designed by Chatterjee *et al*. in the context of a structural characterization study of the NS1 protein [36]. Their work demonstrated the importance of the NS1 α3 helix for IFN antagonism, and showed that the Y125A and L132/133A mutations in this region impair the function of NS1 without affecting its structure. To confirm that reversion to the WT sequence did not occur during viral replication, we performed Sanger sequencing of viral RNA isolated from HNEC apical washes (three transwell inserts from each of three donors per virus, i.e. 9 inserts/virus) and Vero/A549 cell supernatants (one sample per cell line per virus) at 96 hours post infection (hpi). The resulting sequences confirmed that the inserted mutations had not reverted to WT during the first 96 hours of replication in any of the culture systems (**Fig. 1b**). Furthermore, no compensatory mutations were observed in other regions of NS1 or NS2.

**Fig. 1.**
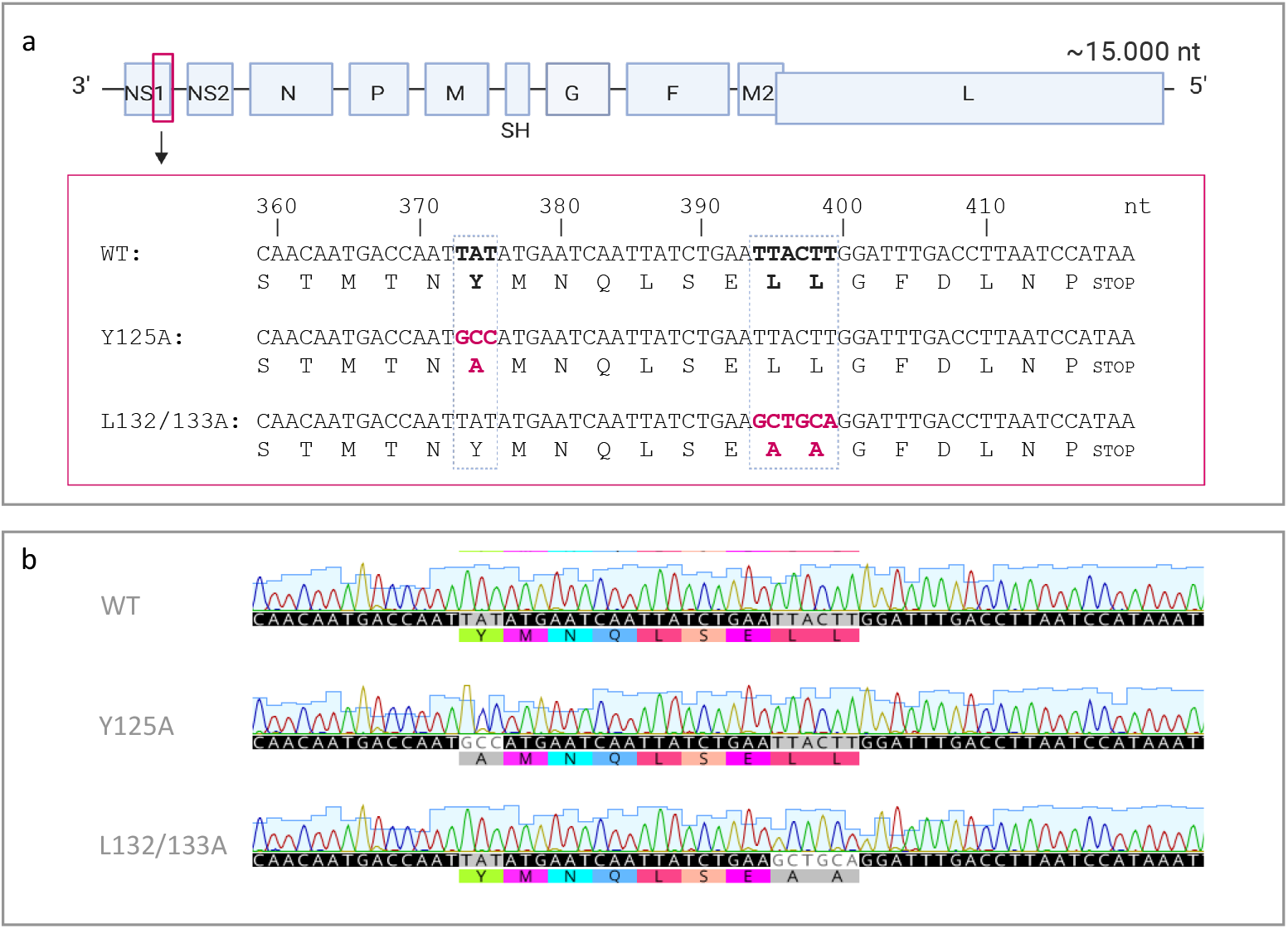
NS1 mutant viruses remain stable during infection of primary HNEC cultures. **(a)** Overview of the RSV genome. Nucleotide substitutions were performed at the indicated positions in the NS1 gene using transformation-associated recombination (TAR) cloning in yeast. **(b)** Sanger sequencing of RNA isolated from HNEC apical wash samples (3 replicates from 3 donors per virus, i.e. 9 samples per virus) and Vero/A549 supernatants (1 replicate per virus) at 96 hpi revealed no reversion or adaptation of the NS1 mutant viruses. Raw sequencing data of 1 HNEC apical wash sample per condition is shown. This data is representative for all other samples. Nucleotide numbering is relative to NS1 gene start.

### RSV NS1 mutants show delayed replication kinetics in HNEC grown at air-liquid interface

It was previously observed by Chatterjee *et al*. that RSV without functional NS1 showed reduced replication in interferon (IFN)-competent cell lines, whereas replication in interferon incompetent cell lines was unaffected [36]. For this reason, we first aimed to reproduce these growth curve experiments on Vero cells (lacking IFN) and A549 cells (IFN competent) **(Fig. 2 a-d)**. The production of viral progeny was quantified using both qPCR and TCID50 assays on Vero cells, to assess both total and infectious virus progeny. In contrast to existing literature [31, 36], our data show that replication of NS1 mutant viruses is similar to WT virus in both Vero and A549 cells **(Fig. 2 a-d)**. We then performed growth curve experiments on HNEC cultured at ALI. Cultures were apically infected, and apical wash samples were obtained at 2, 24, 48, 72 and 96 hpi. In contrast to immortalized cell lines, in primary HNECs, both qPCR and TCID50 assays showed a slight delay in replication of the two NS1 mutant viruses compared to WT at 48 and 72 hpi **(Fig. 2 e,f)**. At 96 hpi all three viruses had reached similar titers.

**Fig. 2.**
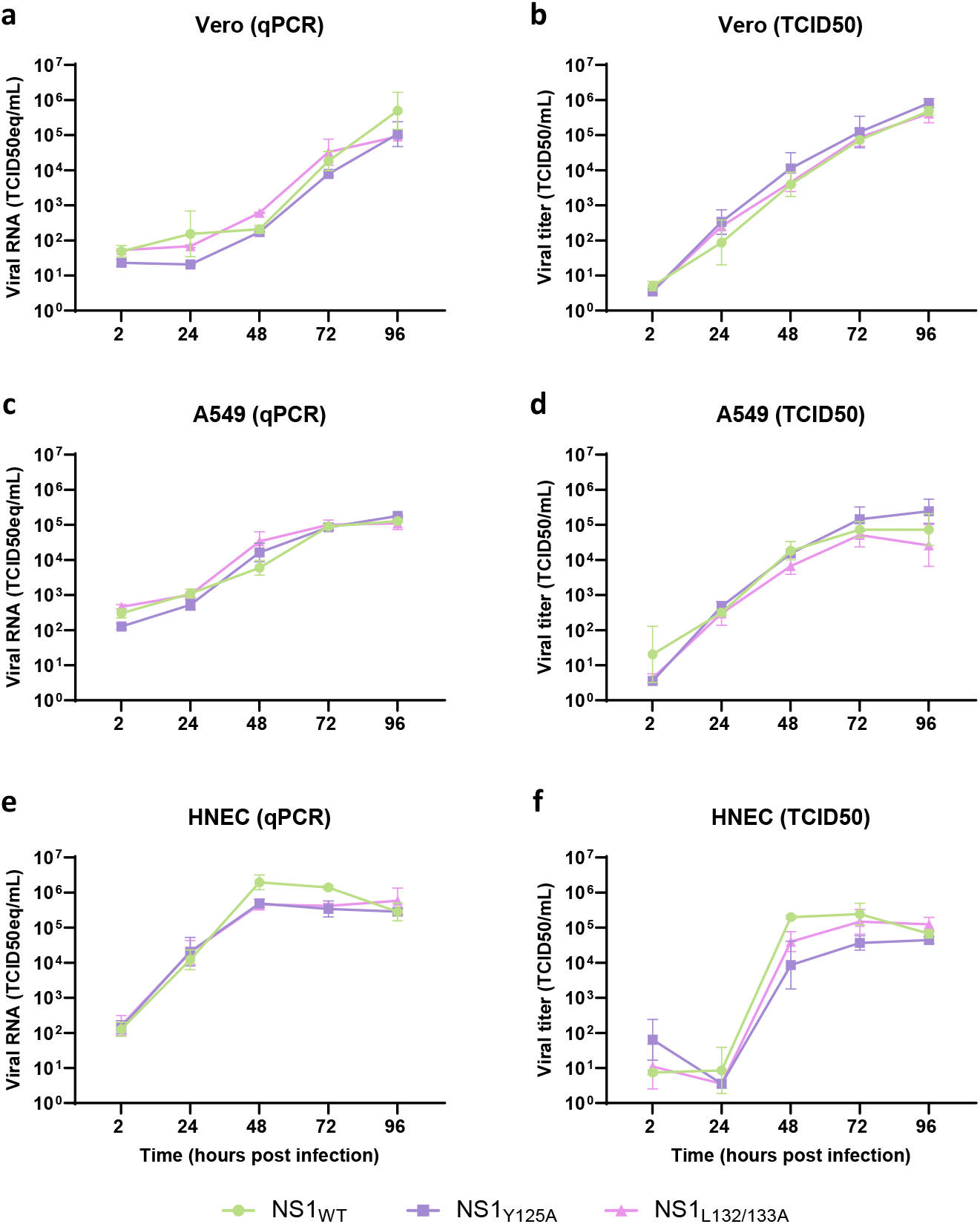
Growth kinetics of WT and NS1 mutant RSV on immortalized cell lines and differentiated primary human nasal epithelial cells (HNEC) cultured at air-liquid interface (ALI). Vero **(a, b)** and A549 **(c, d)** cells were infected in triplicate at an MOI of 0.02. Supernatants were collected at the indicated timepoints. HNEC ALI cultures **(e, f)** from 3 donors were infected in triplicate for each donor using an inoculum of 1×10^5^ TCID50_Vero_ per transwell insert. At the indicated timepoints, apical washes were collected. RT-qPCR was performed on RNA isolated from cell culture supernatant and apical wash samples **(a, c, e)**. Additionally, viral titers were determined by TCID50 assay on Vero cells in quadruplicate technical replicates **(b, d, f)**. Graphs show the geometric mean + geometric SD from 3 technical replicates (Vero/A549) or of 3 biological replicates (mean of 3 transwell inserts for each donor; HNEC).

### RSV NS1 inhibits IFN and chemokine gene expression in primary HNEC ALI cultures

Inhibition of IFNs by RSV NS1 was previously demonstrated in undifferentiated immortalized epithelial cell lines using complete NS1 deletion mutants or ectopic expression of α3 helix mutants [31, 36]. To obtain a more complete picture of the effect of NS1 on host gene expression during infection, we compared the early host response to infection with RSV WT or NS1 mutants in differentiated HNECs cultured at ALI **(Fig. 3a**). To avoid effects due to differences in replication kinetics as much as possible, bulk RNA sequencing was performed at 20 hpi, with mock-infected cultures as control. A differential gene expression (DEG) analysis was performed for both mutant viruses compared to the WT virus (**Fig. 3b** and supplementary dataset). As expected, several interferons (IFN-β1, IFN-λ1/2/3) showed significantly upregulated gene expression levels in both NS1 mutant virus-infected cultures compared to the WT virus (**Fig. 3c**). In addition, the expression of several chemokines (CXCL9, CXCL11, CCL5) was significantly upregulated in cultures infected with either of the NS1 mutant viruses compared to WT virus. In addition to these IFNs and chemokines, there is an upregulation of genes that play a role in the activation of the adaptive immune response, for example TNFSF13b (B-cell activating factor BAFF) and ICOS (inducible T-cell costimulator) in NS1 mutant-infected compared to WT-infected cultures. To provide more context to these data, a gene-set enrichment analysis (GSEA) was performed using the Gene Ontology (GO) database (**Fig. 3d**). A complete list of the identified pathways for both mutant viruses can be found in the supplementary material. Among the identified pathways were ‘regulation of cell-cell adhesion’, ‘response to virus’ and ‘regulation of T cell activation’. Differentially expressed genes belonging to these specific pathways are shown in a heatmap in **Figure 3e**. These data show that the NS1 mutant-infected HNEC similarly display a considerable upregulation of gene expression, whereas gene expression in the WT condition is only slightly upregulated compared to mock.

**Fig. 3.**
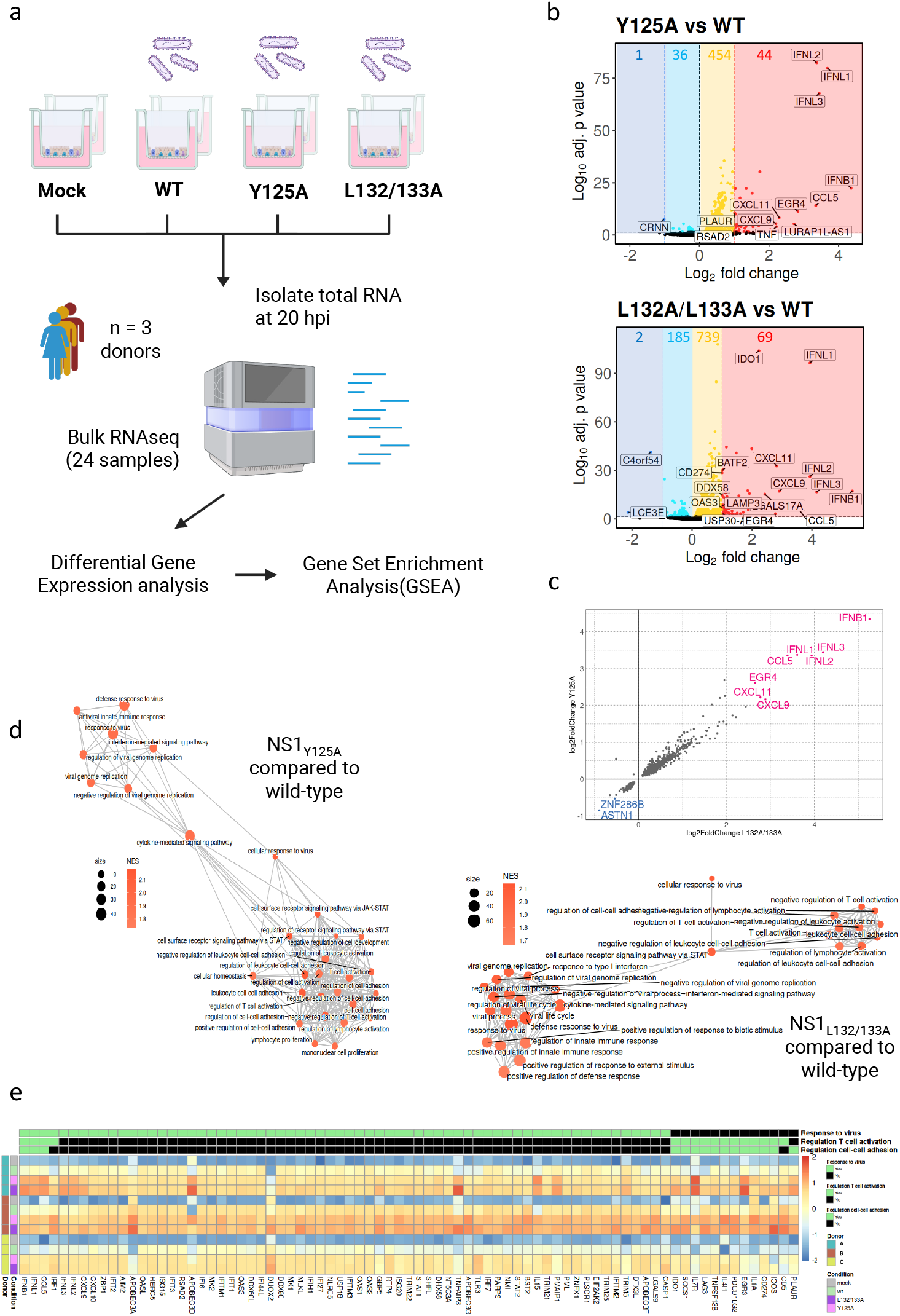
Bulk RNA sequencing of primary HNEC after infection with WT or NS1 mutant RSV. **(a)** Schematic overview of the experimental workflow for bulk RNA sequencing. Differentiated primary human nasal epithelial cells cultured at air-liquid interface were infected with RSV NS1_WT_, NS1_Y125A_ or NS1_L132/133A_ using 1×10^5^ TCID50_Vero_ per transwell or mock-infected. At 20 hpi, RNA was isolated from whole cell lysates and subjected to bulk RNA sequencing. Differential expression gene (DEG) analysis was performed by comparing each mutant to the wild-type condition. **(b)** Volcano plots showing the log fold change of each gene over their adjusted *p* value in log scales. Each dot represents a gene. Colored genes correspond to statistically significant gene expression changes (adjusted *p* value <0.05). Genes in dark blue (logFC <-1) or cyan (logFC ≥-1 and logFC <0) are downregulated. Genes in red (logFC >1) or yellow (logFC ≤1 and logFC >0) are upregulated. **(c)** Correlation plot showing the common significantly altered genes (adjusted p value < 0.05) between both NS1 mutants compared to WT. Common genes with the strongest gene expression changes are highlighted in pink (logFC >2) and blue (logFC <-0.5) **(d)** Enrichment map showing gene ontology (GO) terms enriched from each gene set enrichment analysis (GSEA); edges connect overlapping gene sets. Size of the nodes represent the number of genes included in that GO term and the color represent the normalized enrichment score (NES). **(e)** Heatmap showing the normalized scaled counts of the core enriched genes from GO terms of interest commonly reported in the GSEA for both of the NS1 mutants.

### RSV NS1 mutant viruses induce enhanced levels of IFNs and chemokines in basolateral medium in differentiated HNEC early after infection

After demonstrating that the NS1 mutant viruses induce a more pronounced upregulation of IFN and chemokine gene expression in HNEC compared to wild-type early after infection, we investigated whether this translates to increased secretion of these mediators by the epithelial cells. Cytokine levels were assessed in the basal medium at different timepoints up to 96 hpi, using a Legendplex assay. To visualize the dynamics of cytokine secretion by NS1 mutant-compared to WT-infected cultures, concentrations are depicted as fold change compared to the WT condition for each timepoint in **Figure 4**. Absolute levels for each cytokine can be found in **Figure 5**. In line with the gene expression data shown in **Figure 3**, at early timepoints post infection, increased levels of IFNs and chemokines were present in the basal medium of HNEC that had been infected with RSV NS1 mutant viruses compared to the WT virus (**Fig. 4**). More specifically, the IFN lambda and CXCL10 levels had the highest fold change difference at 24 hpi, whereas the levels of IFNα/β, CXCL11 and CCL5 in NS1 mutant-infected cells differed the most from the WT condition at 48 hpi. Absolute cytokine levels showed the highest concentrations at 48 hpi for the NS1 mutant infected HNECs, after which the cytokine levels in the WT-infected cells surpassed these from 72 hpi onwards (**Fig. 5**). For most cytokines, the secretion pattern is similar between the two NS1 mutants, except for IL-6 and CXCL10, which are mainly secreted upon infection with NS1_Y125A_ and NS1_L132/133A_, respectively.

**Fig. 4.**
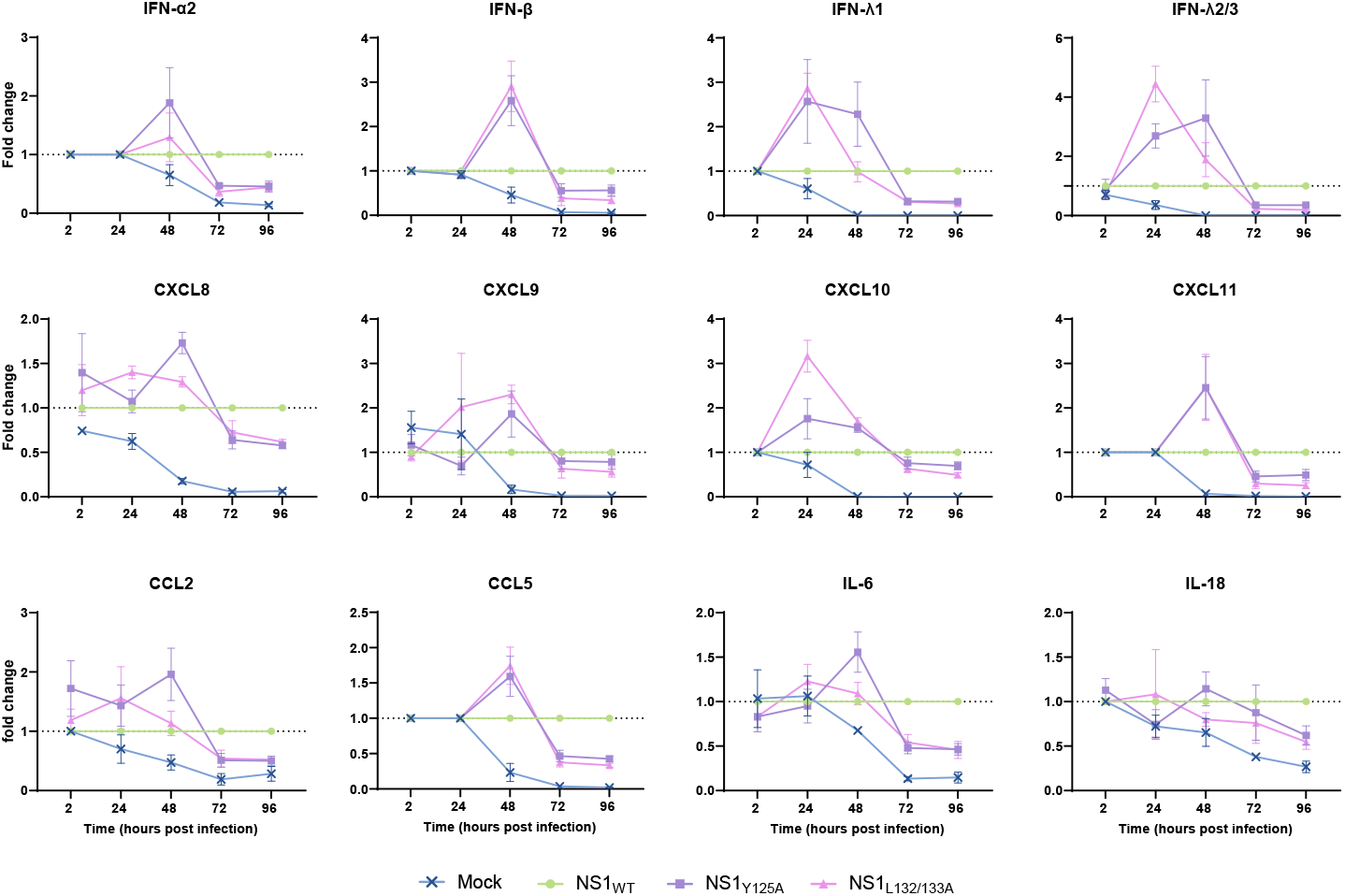
Basolateral cytokine secretion by human nasal epithelial cell (HNEC) cultures grown at air-liquid interface (ALI) infected with WT and NS1 mutant RSV – fold change compared to WT. ALI cultures from 3 donors were infected in triplicate using an inoculum of 1×10^5^ TCID50_Vero_ per transwell insert. At the indicated timepoints, basolateral medium samples were collected. Cytokines were measured using a custom Legendplex panel and fold change was determined with respect to WT levels for each timepoint. All concentrations below the detection limit were set to ([detection limit]/2). Graphs show the mean + SEM from 3 independent epithelial donors.

**Fig. 5.**
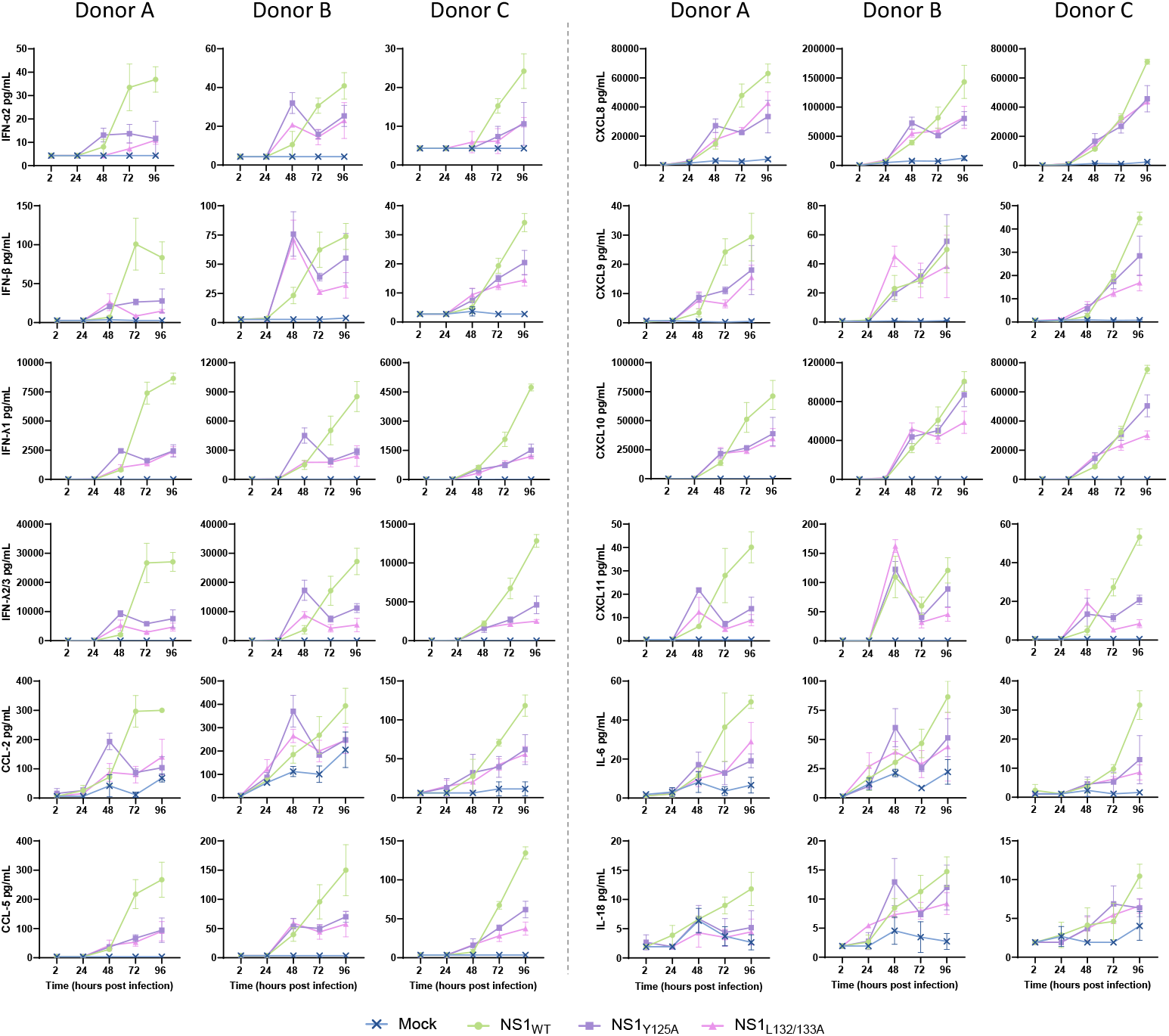
Basolateral cytokine secretion by human nasal epithelial cell (HNEC) cultures grown at air-liquid interface (ALI) infected with WT and NS1 mutant RSV – absolute cytokine levels. ALI cultures from 3 donors were infected in triplicate using an inoculum of 1×10^5^ TCID50_Vero_ per transwell insert. At the indicated timepoints, basolateral medium samples were collected. Cytokines were measured using a custom Legendplex panel. All concentrations below the detection limit were set to ([detection limit]/2). Graphs show the mean + SD from 3 technical replicates for each epithelial donor separately.

### RSV-infected primary HNEC grown at air-liquid interface attract neutrophils

Considering that the attraction of immune cells is an important function of infection-induced chemokines, we assessed whether there are differences in the level of migration induced by NS1 mutant and WT viruses. In order to assess this, we set up an indirect migration assay to determine attraction of whole blood-derived PBMC and granulocyte fractions to medium containing cytokines released by (mock-)infected HNEC cultures (for a schematic overview see **Fig. 6a**). The basal (migrated) fraction was stained with α-CD45, before mixing with the apical fraction (unmigrated, unstained). All cells were stained with a PBMC/granulocyte antibody panel, and analyzed using flow cytometry. For basolateral medium harvested from infected HNEC cultures at 24 hpi, no clear differences in total immune cell migration could be observed between the different experimental conditions. However, for basolateral medium samples harvested at 48 hpi a clear increase in total immune cell migration could be observed towards supernatants from RSV-infected HNECs compared to supernatant from the mock condition **(Fig. 6b)**. In addition, a trend can be seen towards increased immune cell migration in the NS1 mutant conditions compared to the WT condition at 48 hpi. When the immune cell migration was analyzed per immune cell type, it appeared that especially neutrophil migration showed a very similar pattern to the total immune cell migration **(Fig. 6c)**. Indeed, when the migration of all other immune cell types combined (non-neutrophils) was plotted, the trend disappeared **(Fig. 6d)**. This suggests that the increase in total immune cell migration in the WT and NS1 mutant conditions can be mainly explained by neutrophil migration.

**Fig. 6.**
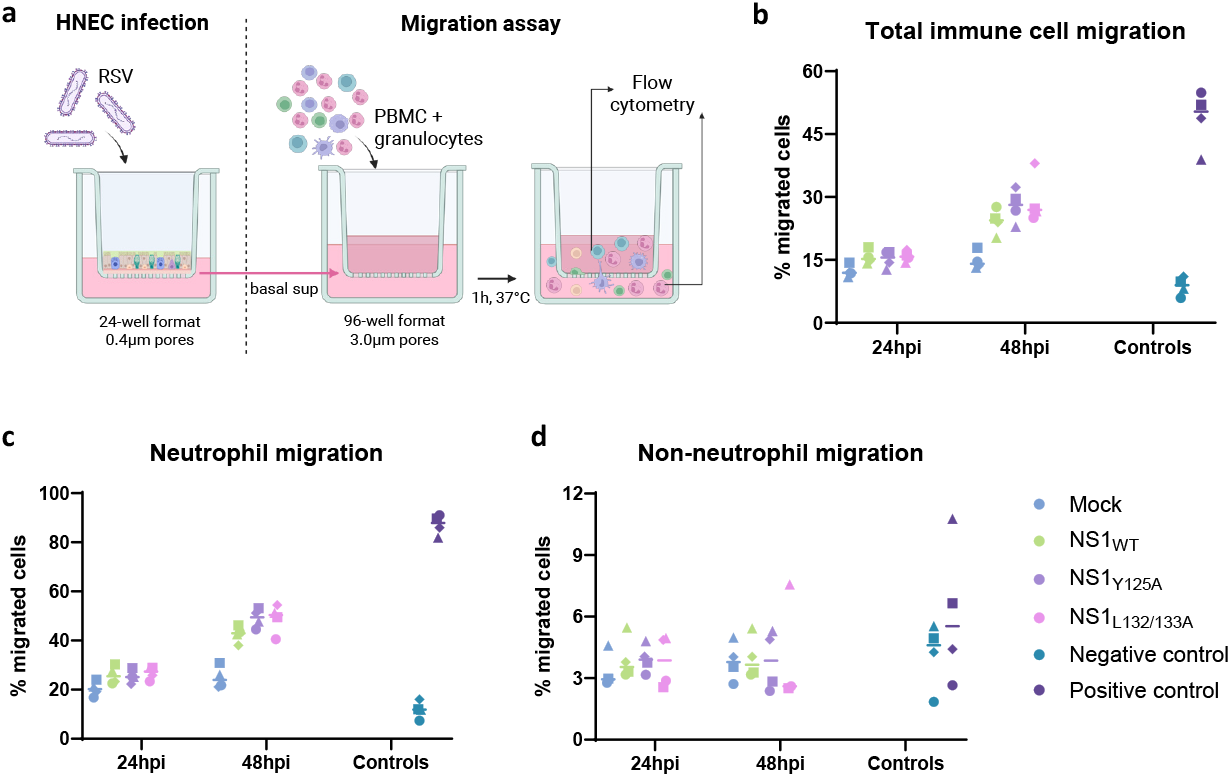
Medium from RSV-infected cultures induces neutrophil migration. **(a)** An indirect immune cell migration assay was performed using the isolated PBMC and granulocyte fractions obtained from fresh whole blood of healthy adults volunteers. Cells were allowed to migrate through 3 μm pore-size membranes for 1 h towards media obtained at 24 and 48 hpi from RSV-infected HNEC cultures, pooled from 3 epithelial donors. Migrated and non-migrated fractions were characterized using flow cytometry. Percentage of migrated **(b)** total PBMC and PMN, **(c)** neutrophils, and **(d)** non-neutrophils (CD66^-^) for media obtained from mock-, WT- and NS1 mutant-infected HNEC cultures. The data was obtained in 2 independent experiments using 4 blood donors. Each symbol represents an individual donor (n = 4), tested in triplicate. Lines represent the median.

## Discussion

Here, we have studied the replication kinetics of RSV WT and NS1_Y125A_ and NS1_L132133A_ mutants, both in immortalized cell lines and in physiologically relevant differentiated primary human nasal cells cultured at air-liquid interface. In Vero and A549 cells, both NS1 mutants showed replication kinetics that were comparable to WT virus. In contrast, a delay in replication of the NS1 mutant viruses compared to WT RSV was observed in HNEC cultured at ALI. Bulk RNA sequencing of these primary nasal cell cultures showed an increased expression of interferons and chemokines in cultures that were infected with NS1 mutants compared to the WT virus in the initial phase of infection. These findings were further confirmed by analysis of the basal media from these cultures, in which increased levels of IFNs and chemokines were detected at early timepoints post infection of HNECs infected with NS1 mutants compared to WT. Finally, we have demonstrated that the basal media from RSV-infected HNECs induce migration of neutrophils.

In contrast to previous studies, we did not observe decreased replication of NS1 mutants compared to WT RSV in immortalized cell lines. Spann *et al*. previously found reduced replication of a complete NS1 gene deletion mutant in both Vero and A549 cells [31]. In light of our observations, this replication defect might be explained by a more general effect of this gene deletion on viral replication, rather than being due to a specific effect on immune evasion. In addition, Chatterjee *et al*. have shown reduced replication of RSV-A2 encoding the NS1_Y125A_ and NS1_L132/133A_ mutations in A549, but not Vero cells [36]. A possible explanation for this observation could lie in the way the virus stocks were produced for their study. As this is not explicitly described in the method section, the viral stocks used in their study might not have been sucrose-purified, which may have resulted in the presence of IFNs and other cytokines that were secreted by the cells in which the virus was produced. These cytokines might have interfered with viral infection and/or replication, especially in an IFN-competent cell line like A549. The virus stocks used for our study have been sucrose-purified, which reduces the content of contaminating cytokines present in the inoculum.

As expected, bulk RNA sequencing data from HNEC cultures at 20 hpi showed an increased expression of IFN-β and IFN-λ upon infection with the NS1 mutants compared to WT virus. This could be an explanation for the delayed viral replication that is observed up to 72 hpi. In addition to the interferons, several chemokines showed an upregulated expression in NS1 mutant-infected cells. This indicates the induction of a stronger immune response, and hints towards recruitment and activation of immune cells at an early stage of the infection. This is further confirmed by increased expression of the B-cell activating factor BAFF and the T cell activator ICOS. Another interesting finding is the upregulation of IDO1 (indoleamine 2, 3-dioxygenase) expression in NS1 mutant-infected HNEC cultures. IDO1 is an enzyme associated with immune suppression and attenuation, and it was found to be upregulated by IFN-λ in the context of influenza virus infection [50]. IDO1 has also been found to cooperate with IFN-γ to reduce RSV replication *in vitro* [51]. Together, these data show that the NS1 protein plays an immunosuppressive role in the early stages of RSV infection, thereby facilitating viral replication.

In line with the results of our gene expression analysis, HNEC cultures infected with the NS1 mutants showed a peak in IFN and chemokine secretion into the basal medium at 24-48 hpi, after which there was a strong reduction in cytokine secretion in comparison with the WT condition. Although absolute cytokine levels did not drop after 48 hpi, which is likely due to the presence of residual cytokines in the basal medium, the fold change in secretion approached that of mock levels at 72-96 hpi. This corresponds well with a study by Pei *et al*., which shows that the gene expression pattern between mock- and NS1_Y125A_-infected A549 cells was very similar at 96 hpi, whereas WT RSV-infected cells showed highly increased expression of various immune-related molecules at this timepoint [52]. The beneficial effect of an early peak in cytokine secretion *in vivo* is demonstrated in a human challenge study by Habibi et al., in which 58 individuals were infected with RSV [53]. Of these, 23 individuals developed PCR-positive RSV infection with cold-like symptoms, whereas the remaining participants did not develop symptoms. Interestingly, the individuals that did not develop a cold, showed an early peak in secretion of cytokines and chemokines like IL-6, CXCL10, CCL5 and CCL3. This suggests that a strong, early cytokine response can indeed prevent the development of a symptomatic RSV infection. In contrast, according to the same study, a delayed peak in cytokine secretion correlates with the development of disease [53].

Chemokines like CXCL8, CXCL10, CXCL11 and CCL5 are known for their ability to attract cells of both the innate and adaptive immune system. The presence of these chemokines has been demonstrated in broncho-alveolar lavage (BAL) and nasal fluids of RSV-infected individuals [54, 55, 24]. In some studies, chemokine levels (or the ratios between them) have been found to correlate with disease severity [56]. The role of immune cell recruitment during RSV is somewhat ambiguous: on the one hand these cells may mediate a protective antiviral response, but on the other hand they have been associated with immunopathology, especially during severe RSV disease [57]. In light of this, it would be beneficial for the infected epithelium to only secrete these chemokines for a brief period, like we observed in the NS1 mutant-infected HNECs, and also much like the observations that were made in the human challenge study mentioned before [53]. This way, an antiviral immune response can be induced, and subsequently stopped before it causes excessive harm to the epithelium. This corresponds with data that was obtained in (naïve) non-human primates, in which infection with an RSV-A2ΔNS1 strain led to reduced viral replication, but was still found to be immunogenic and protective against subsequent RSV-A2 WT infection [18]. Such characteristics are ideal for a live-attenuated vaccine strain. The fact that we can recapitulate these kinetics in our primary HNECs at ALI, demonstrates that this could be a suitable model to perform a preclinical assessment of RSV vaccine candidates and to monitor their effects on the respiratory epithelium. In addition to the characterization of viral replication kinetics, this model can also be used to predict cytokine and chemokine secretion profiles upon RSV infection.

Despite the many benefits of the air-liquid interface respiratory epithelial transwell model, a downside of this model remains the absence of cells from the immune system. Although many attempts have been made, to the best of our knowledge, only a few studies have demonstrated successful direct co-cultures between respiratory epithelial cells at ALI and immune cells [58, 59]. To allow for immune cell migration, 3 μm pores are needed, whereas the HNECs are cultured on 0.4 μm pore-size membranes. Additionally, different cell types would require different media compositions, which could affect both epithelial and immune cell responses. In this study, we therefore employed an indirect approach to study the interaction between the epithelium and various immune cell fractions, by allowing immune cells to migrate towards basal supernatants from infected HNEC in a separate assay. Our data show a clear increase in migration of immune cells towards media from RSV-infected HNEC, both from the WT and NS1 mutant conditions, harvested at 48 hpi. In addition, basal media from NS1 mutant-infected HNEC seem to induce the highest level of immune cell migration. This trend corresponds well with the chemokine levels that were detected in the basal medium of these cultures. Looking at the separate immune cell types, the observed increase in cell migration towards the RSV-infected conditions seems to mainly involve neutrophils. The exact role of neutrophils in RSV infection is still debated. While some studies describe a positive role for these cells in viral clearance, others stress their exacerbating effects on lung pathology, and one study shows that neutrophils do not seem to have an effect on RSV infection at all [60-62]. A complex mix of environmental factors seems to determine whether neutrophil recruitment has a positive or negative impact on viral clearance in the infected tissue [63]. Because our assay does not involve a direct interaction between neutrophils and the epithelium, it is not possible to assess the exact effects of this neutrophil recruitment on the course of infection.

Another limitation of our study is that the adult epithelial antiviral response differs from that in infants, especially those below the age of 1 year, which is the most important risk group for severe RSV disease [64]. The cytokine profiles detected in our primary HNEC model might accurately predict the *in vivo* epithelial response in adults, but they could misrepresent the situation in the developing airways of young children. To assess the infant epithelial immune responses, the use of pediatric epithelial cells would be more relevant. ALI transwell cultures of pediatric epithelial airway cells have been described in several studies, and it seems that their characteristics with respect to the antiviral immune response correspond quite well with the *in vivo* situation [65-67]. This is an important consideration when *in vitro* airway epithelial models are utilized for the assessment of potential RSV vaccine candidates targeting different risk groups.

In conclusion, to the best of our knowledge, this is the first study to describe the effect of RSV NS1 α3 helix mutations in differentiated HNECs cultured at ALI. We have shown that this model closely resembles the *in vivo* situation in both viral replication and in the production of cytokines and chemokines. This makes the HNEC model an excellent tool for the early assessment of live-attenuated RSV vaccine candidates, as viral replication and chemokine data could provide an indication of the strength and direction of a subsequent vaccine-induced immune response. With this, our study provides insights that will help move the field of live-attenuated RSV vaccine development forward.

## Acknowledgements

We thank Sanjay Vashee (JCVI) and Natalay Kouprina (NIH) for kindly providing vector pCC1BAC-His3 and yeast strain VL6-48N, respectively. The following reagents were obtained through BEI Resources, NIAID, NIH: RSV-A2 L/N/P/M2-1 Helper Plasmids (NR-36461/NR-36462/NR-36463/NR-36464).

## Data availability statement

Raw and processed RNAseq data is available in the Gene Expression Omnibus database (GEO, https://www.ncbi.nlm.nih.gov/geo/) with accession number GSE309353.

## Author contribution statement

RWK: Conceptualization, Formal Analysis, Investigation, Writing – Original Draft, Writing – Review & Editing, Visualization; AMG: Formal Analysis, Data Curation, Writing – Review & Editing, Visualization; NE: Methodology, Resources, Writing – Review & Editing; MFL: Methodology, Writing – Review & Editing; AJL: Investigation, Writing – Review & Editing; HJH: Methodology, Investigation, Writing – Review & Editing; ATG: Methodology, Investigation, Writing – Review & Editing; JdJ: Writing – Review & Editing, Project Administration; CACMvE: Writing – Review & Editing, Supervision, Funding Acquisition; VT: Methodology, Resources, Writing – Review & Editing; RD: Methodology, Resources, Writing – Review & Editing; PBvK: Conceptualization, Methodology, Writing – Review & Editing, Supervision, Project Administration, Funding Acquisition All authors reviewed and approved the final manuscript.

