## Supplementary Figure 1 for "Respiratory syncytial virus nonstructural protein 1 inhibits production of cytokines and chemokines by differentiated primary nasal epithelial cells cultured at air-liquid interface"

### Supplementary figure S1

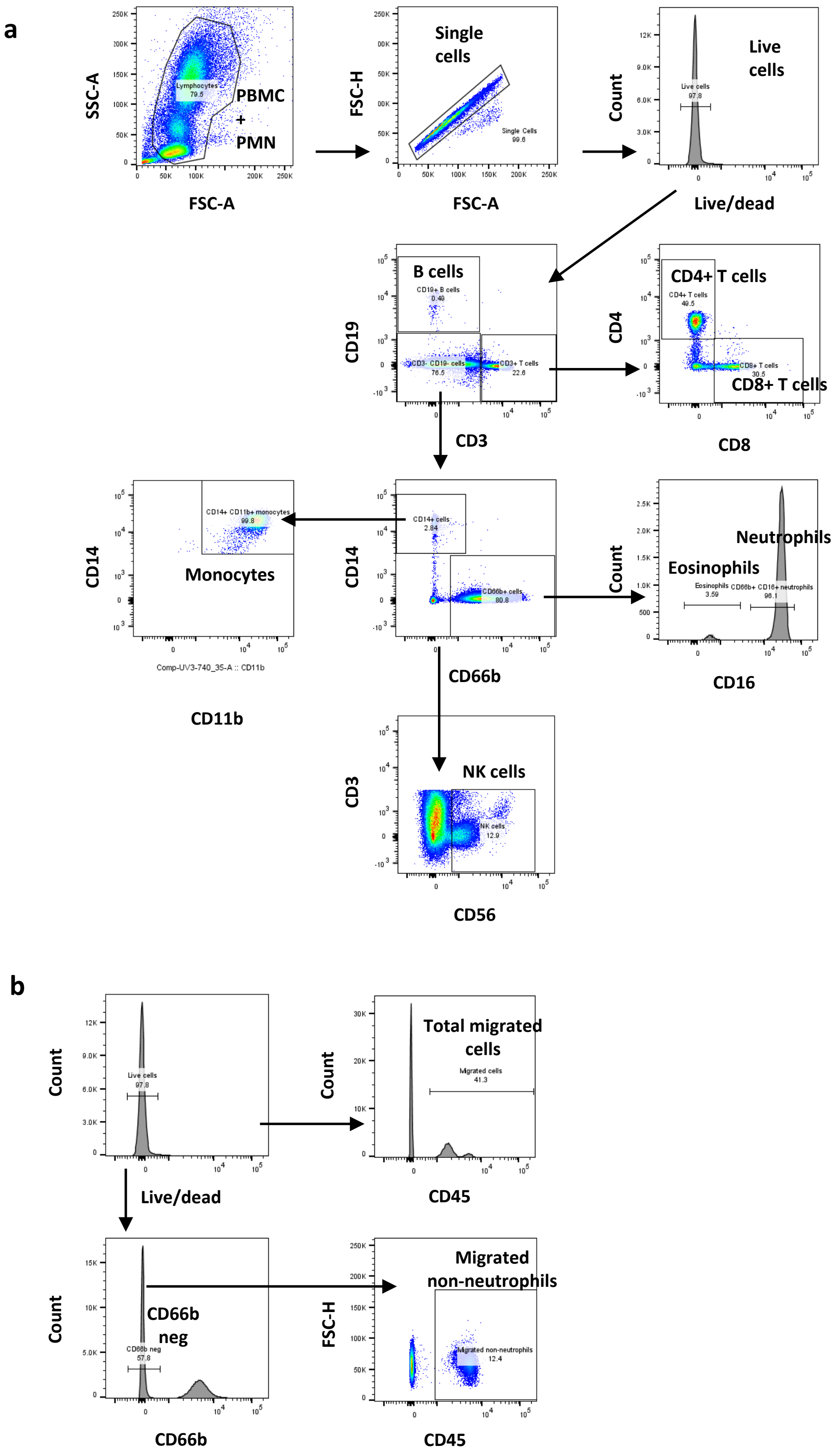


**Fig. S1. Gating strategy for flow cytometry of indirect migration assay. (a)** Gating strategy for individual cell types including neutrophils, eosinophils, NK cells, monocytes, B cells, CD4 and CD8 T cells. **(b)** Gating strategy for total, neutrophil and non-neutrophil migrated cells.
